# Effect of stimulus polarity on detection thresholds in cochlear implant users: relationships with average threshold, gap detection, and rate discrimination

**DOI:** 10.1101/297085

**Authors:** Robert P. Carlyon, Stefano Cosentino, John M. Deeks, Wendy Parkinson, Julie A. Arenberg

**Affiliations:** Medical Research Council Cognition and Brain Sciences Unit, University of Cambridge, 15 Chaucer Road, Cambridge, CB2 7EF, United Kingdom.; Department of Speech and Hearing Sciences, University of Washington 1417 NE 42nd St., Seattle, WA 98105, USA.

**Author notes:** Corresponding author: Robert P. Carlyon, MRC Cognition and Brain Sciences Unit, 15 Chaucer Road, Cambridge, CB2 7EF, UK.

## Abstract

Previous psychophysical and modelling studies suggest that cathodic stimulation by a cochlear implant (CI) may preferentially activate the peripheral processes of the auditory nerve, whereas anodic stimulation may preferentially activate the central axons. Because neural degeneration typically starts with loss of the peripheral processes, lower thresholds for cathodic than for anodic stimulation may indicate good local neural survival. We measured thresholds for 99-pulse-per-second trains of triphasic (TP) pulses where the central high-amplitude phase was either anodic (TP-A) or cathodic (TP-C). Thresholds were obtained in monopolar mode from four or five electrodes and a total of eight ears from subjects implanted with the Advanced Bionics CI. When between–subject differences were removed, there was a modest but significant correlation between the polarity effect (TP-C threshold minus TP-A threshold) and the average of TP-C and TP-A thresholds, consistent with the hypothesis that a large polarity effect corresponds to good neural survival. When data were averaged across electrodes for each subject, relatively low thresholds for TP-C correlated with a high “upper limit” (the pulse rate up to which pitch continues to increase) from a previous study [Cosentino S, Carlyon RP, Deeks JM, Parkinson W, Bierer JA (2016) Rate discrimination, gap detection and ranking of temporal pitch in cochlear implant users. J Assoc Otolaryngol 17:371– 382]. Overall the results provide modest indirect support for the hypothesis that the polarity effect provides an estimate of local neural survival.

## Introduction

Many cochlear implant (CI) listeners understand speech well, at least in quiet. However there remains substantial across-listener variability, with some struggling even in favorable listening conditions (Holden et al., 2013). In recent years there has been considerable interest in identifying the reasons for poor speech perception, and in identifying the relationship between performance and the global and local pattern of neural survival in each individual patient. A potentially useful approach comes from single-electrode psychophysical measures, which have revealed substantial across-listener and across-electrode variability in a number of tasks. These include signal detection in quiet and supra-threshold measures. Authors have investigated the variation in not only the absolute level of thresholds (Pfingst and Xu, 2004; Bierer et al., 2015), but also how they are influenced by pulse rate (“multi-pulse integration, MPI”; Zhou and Pfingst, 2014; Zhou et al., 2015; Zhou and Pfingst, 2016), pulse polarity (Macherey et al., 2017), and stimulation mode (Bierer, 2007; Bierer and Faulkner, 2010). Supra-threshold tasks have included modulation detection (Garadat et al., 2012; Garadat et al., 2013), gap detection (Bierer et al., 2015), and rate discrimination both at low and high rates (Cosentino et al., 2016). It is worth noting that variation in all of these measures has been observed even when stimulating in monopolar mode, which is believed to produce a wide spread of excitation within the cochlea. Indeed, even though the variation in thresholds across the electrode array is smaller in dB for monopolar than for tripolar stimulation, this is not true when the across-electrode standard deviation (s.d.) is normalized by the within-electrode s.d. (Bierer et al., 2015). In other words, monopolar stimulation may reveal across-electrode variation as reliably as tripolar stimulation, even though the size of this variation is smaller in monopolar than in tripolar mode when expressed in dB.

A potentially important application of single-electrode measures is that they may guide the clinician in choosing which, if any, electrodes to de-activate when optimizing patient maps. Indeed, significant improvements in speech perception scores have been obtained by de-activating electrodes based on high modulation detection thresholds (Garadat et al., 2012; Garadat et al., 2013) and high thresholds for low-rate pulse trains (Zhou, 2017). In order to provide a principled approach to channel selection it would be useful to know how the various different single-electrode measures correlate with each other. This could then either reveal clusters of tests, each of which tap a particular consequence of neural degeneration, or reveal a single test factor that could be used to guide channel selection algorithms. This information may also provide basic insights into the limitations of hearing by CI users. For example, Cosentino *et al*. (2016) found that the “upper limit” of temporal pitch – defined as the highest pulse rate on a single electrode above which pitch no longer increased – correlated significantly with gap detection thresholds (GDTs), but not with the smallest difference in the rate of a low-rate pulse train that could be discriminated. The significant difference between these two correlations led them to suggest that there is a limitation specific to tasks that require sustained temporally accurate firing to high pulse rates, and which is separate from that which limits low-rate discrimination. Zhou and colleagues (2016) reported that MPI correlated significantly with the degree of spatial selectivity for a given electrode, and concluded that integration of multiple pulses is most efficient when conveyed by neurons that innervate a wide region of the cochlea.

The present study forms part of a series that compares performance on different single-electrode psychophysical measures in a group of CI users (Bierer et al., 2015; Cosentino et al., 2016). Here we measure polarity sensitivity, defined as the difference between thresholds for 99-pps trains of triphasic pulses in which the short high-amplitude portion is either anodic or cathodic. As with other types of asymmetric pulse, triphasic stimulation allows one to study polarity sensitivity by concentrating charge of one polarity into a short time period, whilst maintaining the charge balancing necessary for patient safety. All stimulation is in monopolar mode. The motivation stems from the finding that, although animal studies usually reveal greater sensitivity to cathodic than to anodic stimulation (Hartmann et al., 1984; Miller et al., 1999; Miller et al., 2004), the reverse is true for human CI users when presented with stimuli at or close to their most comfortable listening level ("MCL": Macherey et al., 2006; Macherey et al., 2008; van Wieringen et al., 2008; Undurraga et al., 2010; Macherey et al., 2011). A possible reason for this discrepancy, consistent with computational models (Rattay, 1999; Rattay et al., 2001), is that cathodic stimulation depolarizes the peripheral processes of the auditory nerve. These processes are likely to be intact in the recently-deafened animals used in most physiological experiments. However, there is evidence that peripheral processes are more susceptible than central axons to auditory deprivation, and so may have deteriorated in human CI users who have been deaf for months or years prior to implantation (Johnsson et al., 1981).

Polarity sensitivity differs across electrodes not only at MCL but also at threshold. Unlike MCL measures, the direction of the polarity sensitivity at threshold varies consistently across listeners and electrodes, and some electrode-listener combinations reveal lower thresholds for cathodic than for anodic stimulation (Macherey et al., 2017; Mesnildrey et al., 2017). These combinations may reflect local regions of good neural survival in which a relatively high proportion of peripheral processes remain. Here we investigate whether this putative measure of neural survival correlates with measures of gap detection, low-rate discrimination, and the upper limit of temporal pitch obtained in ou previous studies (Bierer et al., 2015; Cosentino et al., 2016). The hypothesis is that lower thresholds for cathodic than for anodic stimulation will correlate with tasks that depend on good local neural survival. Note, however, that this does not require that the variation in performance on those tasks is limited by the pattern of activity in the auditory nerve; rather, poor auditory nerve survival may lead to more central degeneration which in turn could limit performance on perceptual tasks. For example, Carlyon and Deeks (2015) found that the “alternating amplitude” pattern of auditory-nerve evoked responses (Wilson, 1997) to high-rate pulse trains correlated across subjects with poor rate discrimination at high rates, but also demonstrated that the correlation was not causal, and that manipulating the stimulus so as to reduce the alternating-amplitude pattern did not improve performance.

A second prediction is that polarity sensitivity will correlate with the average of the thresholds in the two polarities, on the assumption that better sensitivity to cathodal stimulation will reflect better neural survival and hence lower overall thresholds. Specifically, we assume that the average thresholds will also depend on the distance of the electrodes from the modiolus (electrode-modiolus distance, “EMD”) but that the effects of EMD and polarity are independent. In the discussion section we describe a recent study that provides evidence for this assumption.

When comparing performance on different tasks, two types of measure are possible. One of these is to correlate performance across subjects (e.g., Fu, 2002; Won et al., 2011; Cosentino et al., 2016). This can harness the often substantial across-listener variability in performance, but is potentially susceptible to non-specific effects such as attention span and cognitive ability. Such effects could lead to a correlation that does not reflect any common processing of the two tasks, except at very central levels. A more rigorous approach is to partial out between-subject effects, and to correlate the relative pattern of scores across electrodes (Bierer, 2007; Cosentino et al., 2015; Zhou and Pfingst, 2016). This approach is immune to between-listener cognitive differences. Across-electrode differences are also of more clinical relevance because, as mentioned above, they may guide channel-selection methods that aim to optimize performance on a listener-by-listener basis. However, because this type of analysis excludes the substantial variation in neural survival that occurs across listeners, for example due to differences in pathology (e.g. Zimmermann et al., 1995; e.g. Nadol, 1997), it risks “throwing the baby out with the bath water”. We perform both types of analysis here. One advantage of studying polarity sensitivity is that, being a difference between two thresholds, is unlikely to be affected by between-listener variation in cognition. Hence it may exploit the benefits of measuring the substantial across-listener variation in neural survival without being strongly influenced by cognitive effects.

## Methods

### Subjects

Eight ears from seven post-lingually deafened adults wearing the Advanced Bionics HiRes90K CI were studied; listener details are shown in Table 1. One subject, who was bilaterally implanted, was tested in each ear and is listed as S30L and S39R in the table. This subject’s two ears were treated as completely separate, and therefore, for the purposes of analysis and for discussion in the remainder of this article, there were eight “subjects”. They had all participated in the study by Cosentino *et al* (2016) and the same subject codes are used here. Five of the subjects were implanted and tested in Cambridge, UK, whereas the other three were implanted and tested in Seattle, WA, USA. All procedures were approved by the respective Human Subjects Review Boards.

**Table 1:** Details of the listeners who took part. IDs starting with the letter S refer to patients implanted and tested in Seattle, USA. Those beginning with the letter C were implanted and tested in Cambridge, UK. “Subjects” S30L and S39R refer to the left and right ears of the same participant, and are treated separately throughout this article.

| **ID** | **Age**<br>**(years)** | **Deafness**<br>**Onset (age,**<br>**years)** | **Possible**<br>**Aetiology** | **Duration**<br>**of CI use** |
| --- | --- | --- | --- | --- |
| S22 | 74 | 55 | Hereditary | 7 yr |
| S30L | 51 | 16 | Hereditary | 11 yr |
| S39R | 51 | 16 | Hereditary | 31 yr |
| C1 | 69 | 32 | Unknown | 5 yr |
| C3 | 71 | 50 | Otosclerosis | 4 yr |
| C4 | 68 | 37 | Otosclerosis | 6 yr |
| C5 | 55 | 31 | Unknown | 6 yr |
| C6 | 66 | 51 | Unknown | 3 yr |

### Stimuli

Triphasic pulses were used, in which the amplitude of the central phase was twice that of each of the first and third phases. They are named here in terms of the central high-amplitude phase, which was anodic for stimulus TP-A and cathodic for stimulus TP-C. (Note that here we use the abbreviation “TP” to refer to the triphasic pulse shape, rather than to tripolar stimulation mode as in some other articles). Phase durations were 43 µs, the pulse rate was 99 pulses per second, and the duration of each pulse train was 400 ms. Each subject was tested on the same four or five electrodes as used for the rate discrimination and gap detection measures by Cosentino *et al (2016)*. All stimuli were presented in monopolar mode. Initially MCLs were obtained for each combination of subject and electrode, so as to guide the starting level for the threshold measurements and to set a safety limit for those procedures. After each stimulus was presented the subject indicated its loudness using the Advanced Bionics loudness rating scale for which a “6” is “Most Comfortable” and a “7” is “Loud but Comfortable”. The level was increased gradually until the rating was “7” and then reduced until the listener reported a “6” again. All stimuli were presented and controlled using research hardware and software (“BEDCS”) provided by the Advanced Bionics company. Programs were written using the MATLAB programming environment, which controlled low-level BEDCS routines. Stimuli were checked using a test implant and digital storage oscilloscope. The same software and type of hardware were used at both testing sites (Cambridge and Seattle).

Signal detection thresholds were measured using a three-down, one-up, two-interval forced choice adaptive procedure that converged on 79% correct. Step size was 1 dB for the first two turnpoints and 0.25 dB thereafter. The mean of the last four of six turnpoints were used to estimate threshold. Four repetitions were performed for each measurement. Subjects were asked “Which interval contained the sound?” and responded by selecting a button on a computer screen. Correct-answer feedback was provided at the end of each trial.

## Results

Thresholds (Ts) for each subject are shown for the TP-A (red triangles) and TP-C (blue circles) stimuli in Fig. 1. It can be seen that the polarity effect (T_TP-c_-T_TP-A_) varies across subjects and electrodes. Of the 33 subject-electrode combinations, 9 showed a lower threshold for the TP-C than for the TP-A stimulus, and hence a negative polarity effect. Recall that, according to our hypothesis, low or negative polarity effects reflect good neural survival.

**Figure 1:**
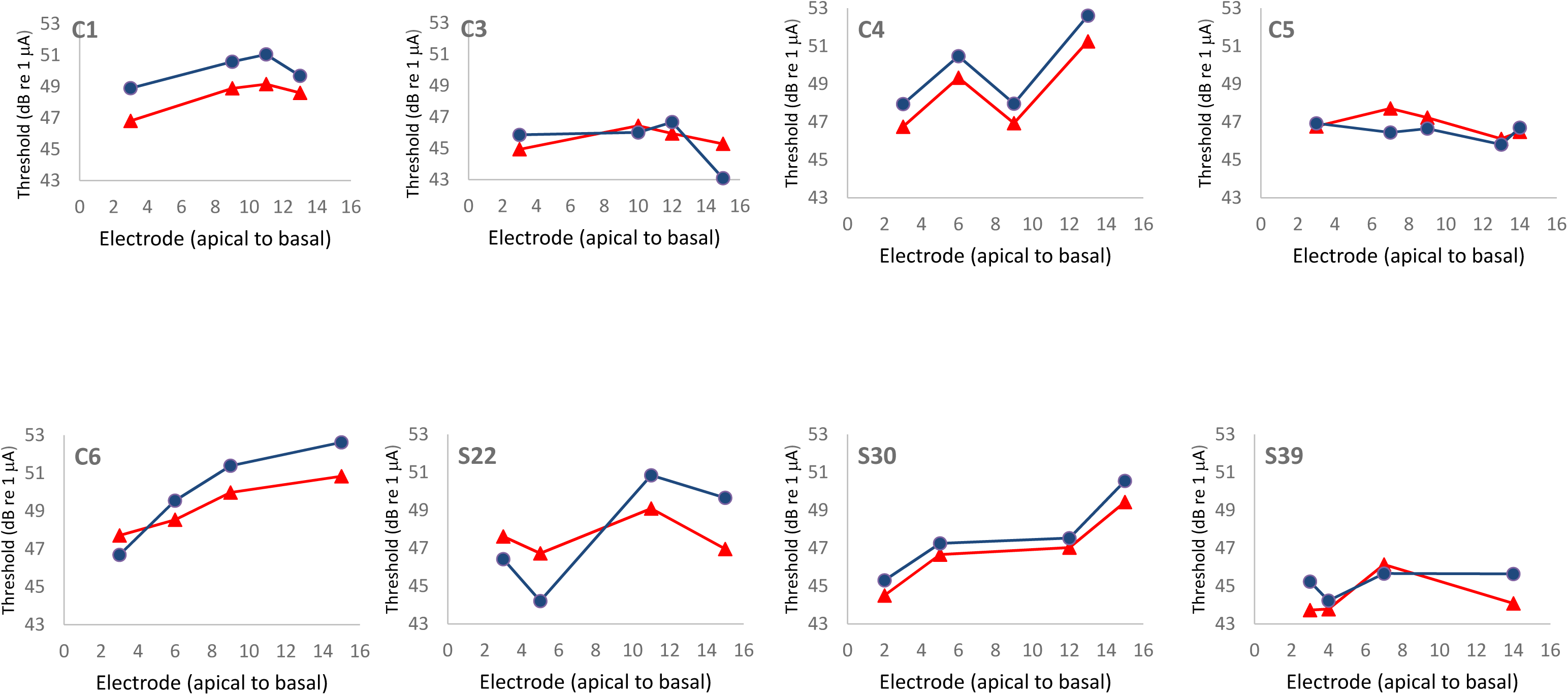
Thresholds as a function of electrode number for each listener. Thresholds for TP-A stimuli are shown by red triangles, whereas those for TP-C stimuli are shown by blue circles.

### Between-electrode correlations

To assess between-electrode correlations we performed univariate ANOVAs with one factor as the dependent variable, the other as a covariate, and with subject entered as a random effect. This revealed a significant positive correlation between the polarity effect and the average of the TP-A and TP-C thresholds (r=0.49, F(1,24)=7.59, p=0.011). Our statistical approach is mathematically equivalent to subtracting each subject’s mean score from every data point, so as to obtain a normalized score, and then correlating these normalized scores. The resulting correlation is shown in Fig. 2. It is in the direction by the hypothesis that lower/negative polarity effects and lower overall thresholds both reflect better survival of the peripheral processes. Note that, although the difference between two scores will *a priori* be correlated with each score alone, no such *a priori* relationship holds between the difference and the *average* of two scores (Oldham, 1962; Tu and Gilthorpe, 2007). This is true even when the two scores correlate with each other. For example, if EMD and neural survival combine additively to affect thresholds, then the across-electrode variation in EMD will produce a correlation between TP-A and TP-C thresholds; however this additive effect will not cause the mean and difference to correlate with each other.

**Figure 2:**
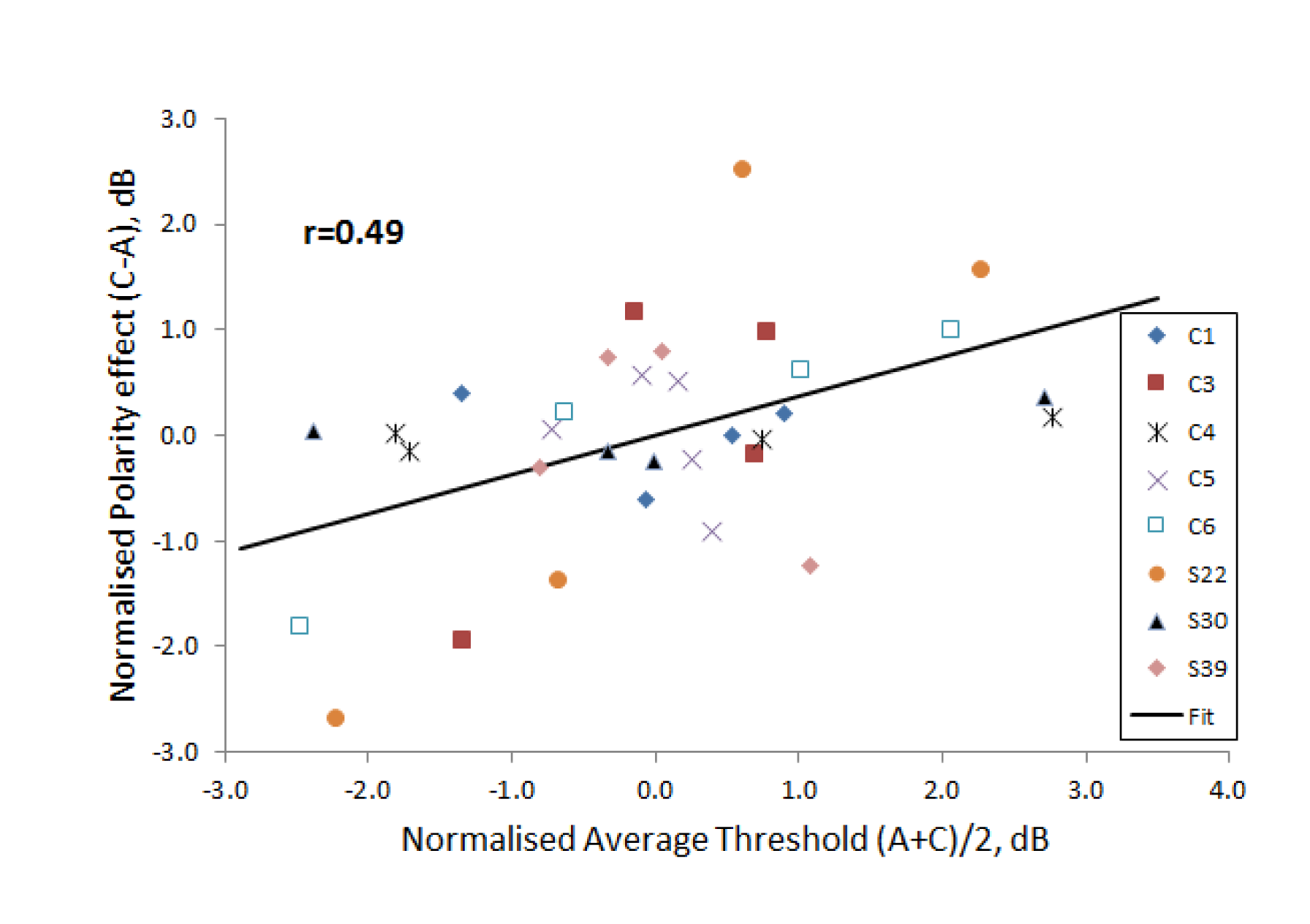
Correlation between the average thresholds (A+C/2) and the difference between cathodic and anodic thresholds (C-A), in decibels. Between subject differences have been removed by subtracting the average value for each subject from every data point for that subject. This normalization removes the effects of between-subject correlations, leaving only the between-electrode correlation. Data for each subject are shown by a unique symbol.

Although there was a significant across-electrode correlation between the polarity effect and the average thresholds, there was no significant correlation between either of them and any of the supra-threshold measures obtained in our previous studies. Those measures were the log of the gap detection thresholds (GDTs) measured by Bierer *et al* (2015), and two rate discrimination measures (“RDR100” and “RDR400”) obtained by Cosentino *et al* (2016). One of those, RDR100, is the log of the ratio between two just-discriminable pulse rates, where the standard had a rate of 100 pps and the signal, whose rate was always above 100 pps was adaptively varied to obtain threshold. The other, “RDR400”, was similar except that the standard had a rate of 400 pps and the signal always had a rate lower than 400 pps. The two measures reflect low-rate discrimination and the “upper limit” of temporal pitch, respectively. In both cases low values represent better performance. The between-electrode correlations are shown in Table 2.

**Table 2:** Across-electrode correlations between the average threshold across polarities and the polarity effect with the logarithms of gap detection thresholds (GDTs), and of rate discrimination measures at low (RDR100) and high (RDR400) rates. The correlations were obtained by subtracting each subject’s mean score from every point, thereby removing between-subject effects. This is equivalent to performing an analysis of covariance with one measure as the dependent variable, the other as a covariate, and subject as a random factor (Bland and Altman, 1995). In all cases there were 24 degrees of freedom. The 1% and 5% significance levels for df=24 are, respectively, 0.50 and 0.39.

|  | GDT | RDR100 | RDR400 |
| --- | --- | --- | --- |
| average | 0.24 | 0.09 | -0.35 |
| polarity | -0.08 | 0.06 | -0.29 |

We also investigated the effect of the longitudinal electrode position in the array. Bierer *et al (2007)* reported a trend for thresholds to decrease from the base to the apex of the electrode array for symmetric biphasic pulses presented in partial tripolar mode, but not in bipolar or monopolar mode. They suggested that this could be due to the progression of spiral ganglion degeneration from base to apex in several pathologies (Zimmermann et al., 1995), and/or to a variation in the EMD along the array. Subsequently, Bierer *et al (2015)* found that thresholds decreased from base to apex for symmetric biphasic pulses presented both in partial tripolar and monopolar mode. A similar finding was observed in the present data for the average of the TP-A and TP-C thresholds, as assessed by a univariate ANOVA with average threshold as dependent variable, subject as a random effect, and electrode number as covariate (F(1,24)=11.41, p=0.002). However, no such finding was observed for the polarity effect (F(1,24)=2.76, p=0.16). Hence, if the polarity effect reflects neural survival, the present results provide no evidence that, for this group of subjects, the effect of longitudinal electrode position on average thresholds was due to a variation in the pattern of neural survival.

### Between-subject correlations

Between-subject correlations were obtained by calculating the average score across electrodes for each subject and then performing product-moment correlations. The correlation between the polarity effect and the average threshold was in the same direction as for the between-electrode correlations, but did not reach statistical significance (r=0.60, df=6, p=0.16). The correlations between each of these measures and the supra-threshold scores obtained previously (GDT, RDR100, and RDR400) are shown in Table 3. All of the correlations were positive, three reached statistical significance, and two did so after Bonferroni correction for multiple comparisons. One of these was between the average threshold and GDT (r=0.87, df=6, p=0.0050, corrected p=0.03). The other was between the polarity effect and RDR400 (r=0.91, df=6, p=0.0017, corrected p=0.01). As noted in the Introduction it is unlikely that the second of these can be explained in terms of cognitive factors, such as the ability or willingness to concentrate on the task, because the polarity effect is a difference score.

**Table 3:** Across-subject correlations between the average threshold across polarities and the polarity effect with the logarithms of gap detection thresholds (GDTs), and of rate discrimination measures at low (RDR100) and high (RDR400) rates. The correlation between the average threshold and the polarity effect was 0.6. Statistically significant correlations are shown in bold; those that survived Bonferonni corrections for six comparisons are additionally underlined (see text for details). The number of degrees of freedom was 6 in all cases. The 1% and 5% significance levels for df=6 are, respectively, 0.83 and 0.71.

|  | GDT | RDR100 | RDR400 |
| --- | --- | --- | --- |
| average | **<u>0.87</u>** | **0.72** | 0.61 |
| polarity | 0.41 | 0.49 | **<u>0.91</u>** |

## Discussion

### Comparison to previous studies

A large number of psychophysical and electrophysiological studies have demonstrated CI users’ greater sensitivity to anodic than to cathodic current for moderately and comfortably loud stimuli (Macherey et al., 2006; Macherey et al., 2008; van Wieringen et al., 2008; Undurraga et al., 2010; Macherey et al., 2011). More recently, two studies have shown that, although detection thresholds do not differ overall between stimuli of opposite polarity, individual CI listeners show idiosyncratic but consistent polarity effects.

Macherey *et al*. (2017) measured thresholds for trains of symmetric biphasic pulses with either anodic or cathodic leading polarity (“SYM-A” and “SYM-C”), as well as for trains of so-called quadraphasic (“QP”) pulses. The QP pulses were constructed by abutting two SYM pulses of opposite leading polarity, such that the central portion consisted of two anodic (QP-A) or cathodic (QP-C) phases. A previous study had shown that, as for the TP-A and TP-C pulse trains used here and for pseudomonophasic pulse trains, the loudness of supra-threshold QP-A pulse trains was consistently greater than that of QP-C pulse trains at the same current (Carlyon et al., 2013). Macherey *et al* (2017) found that, across subjects, the difference between QP-A and QP-C thresholds correlated strongly with that between SYM-A and SYM-C thresholds, demonstrating that the idiosyncratic differences observed across subjects were not simply due to measurement noise. In addition, they reported that, for some electrode-listener combinations, the loudness growth function was non-monotonic, and that this unusual pattern was significantly more likely to occur when thresholds were lower for QP-C than for QP-A pulse trains. They suggested, as we do here, that low QP-C thresholds may reflect good local survival and activation of peripheral processes. They further suggested that nonmonotonic loudness growth might then be explained by “cathodal block” (Ranck, 1975) impeding the propagation of action potentials as level was increased. Interestingly, they only observed non-monotonic growth for users of the Cochlear CI; when they tested five Advanced Bionics users on one electrode each, the loudness growth was always monotonic. The fact that we reported several instances of negative polarity effects raises the possibility that, for those electrode-subject combinations, the loudness growth may also have been non-monotonic. A stronger test of this hypothesis would involve measuring loudness growth functions for the subject-electrode combinations showing the most negative polarity effects, such as C3 (electrode 15, -2.2 dB), C5 (electrode 7, -1.3 dB), and S22 (electrode 5, -2.45 dB).

More recently, Mesnildrey *et al* (2017) measured thresholds using a fast one-interval adjustment procedure for trains of TP-A and TP-C pulses presented in partial tripolar mode. Their measurements, obtained with multiple electrodes in 16 ears, revealed that thresholds were lower for TP-C than for TP-A pulse trains in 22% (48/219) of electrodes. This percentage was very similar to the corresponding value of 27% (9/33) in the present study. Another similarity is that they found a positive across-electrode correlation between the polarity effect and the threshold for a SYM-C pulse train, analogous to our correlation between the polarity effect and overall threshold. However, their correlation was quite weak, accounting only for about 5% of the variance. More recently, Goehring *et al* (2018) used a method similar to that of Mesnildrey *et al* (2017) to measure TP-A and TP-C thresholds in eight subjects. They also reported a weak positive correlation which, in their case, was not significant. Clearly, then, the size and significance of the correlation depends on the particular group of subjects and/or electrodes studied. Indeed, inspection of Fig. 2 reveals that the correlation was, for example, strong for subjects S22 and C6 and absent for others, such as C4 and C5. In terms of our hypothesis, it may be that large correlations occur when the EMD is constant, and neural survival varies markedly, along the length of the cochlea. More generally, it is worth noting that although the across-electrode comparisons reported here and elsewhere involve a reasonably high number of degrees of freedom, they are obtained from only a modest number of subjects.

With the above caveats, of particular interest in the study by Mesnildrey *et al* (2017) is their analysis of CT scans from nine ears, which allowed them to derive an estimate of the electrode-modiolar distance (EMD) for each subject. Although the EMD accounted for between 63 and 68% of the between-subject variance in thresholds, it accounted for only a small proportion of the between-electrode variance. The authors suggested that this was due to EMD varying substantially between but not within ears. They also showed that the EMD did not correlate significantly with the polarity effect. This is important for the interpretation of the present results because it is possible that the site of excitation, and hence the polarity effect, was affected by the EMD. If such a correlation had been found, this would have provided an alternative to our interpretation that the correlation between the polarity effect and overall thresholds was mediated by neural survival.

Zhou and Pfingst (2016) also obtained thresholds for multiple electrodes and estimated EMDs from the data relating EMD to electrode number reported by Long *et al*. (2014). They did not report the correlation directly but did state that MPI, defined as the difference between thresholds for 80- and 640-pps pulse trains, correlated with the EMD. Their data showed generally greater across-electrode threshold variations at 80 pps than at 640 pps, and so it is likely that the variation in the MPI was driven mainly by the 80-pps thresholds. If so, then those thresholds would have correlated positively with the EMD. However, they do not state how much of the variation in 80-pps thresholds was accounted for by EMD, nor whether a substantial across-electrode (rather than across-listener) correlation was observed. In addition it is possible, as suggested by Mesnildrey (2017), that the across-electrode variation in EMD for a given subject may be smaller for Advanced Bionics listeners (as tested here) than for the Cochlear CI24RE listeners studied by Zhou and Pfingst (2016) and by Long *et al* (Long et al., 2014; Zhou and Pfingst, 2016). Mesnildrey reported that the average range of EMDs, across listeners, was 0.75 mm in his study compared to 1.2 mm in Long *et al*’s study; an intermediate value of 1.00 mm is revealed by an analysis of the 10 Advanced Bionics listeners studied by de Vries *et al*. (2016).

### Relationship to supra-threshold tasks

Table 3 shows the across-subject correlations between the two measures obtained here – average thresholds and the polarity effect – and the supra-threshold measures reported previously by our laboratories (Bierer et al., 2015; Cosentino et al., 2016). Two correlations were statistically significant after Bonferroni correction: average thresholds vs. log GDTs, and the polarity effect vs RDR400 (which is a measure of the upper limit of temporal pitch). This does not mean, though, that log GDTs are more strongly related to average thresholds than to polarity, or that RDR400 is more strongly correlated with the polarity effect than with average thresholds. Such a conclusion would require that these two correlations were significantly larger than, respectively, those between polarity vs log GDT and average threshold vs. log400R. This was not the case (Williams test, df=5, two-tailed p=0.07 and 0.14 for GDT and RDR400 respectively). The safest conclusion is that both low average thresholds and a small or negative polarity effect correlate with good performance on some supra-threshold tasks, including gap detection and a measure (RDR400) of the upper limit of temporal pitch. There is also evidence from Mesnildrey *et al* (2017) that, across subjects, small or negative polarity effects correspond to low thresholds on a spectro-temporal ripple test (Aronoff and Landsberger, 2013). In addition, Zhou and Pfingst (2016) showed that high thresholds for 80-pps pulse trains – similar to the rate used here, correlated significantly, both across subjects and electrodes, with poor spatial selectivity as measured using forward masking. A *caveat* is that they measured spatial selectivity by measuring the slope relating masker position to the amount of masking of a fixed probe. This measures the spread of excitation produced by the maskers rather than by the signal.

Finally, it is worth noting that Zhou (2017) has recently observed substantial and significant improvements, both in speech perception and on a spectro-temporal ripple test, by de-activating electrodes with high low-rate thresholds. As they have pointed out, such high thresholds may reflect a combination of large EMD and poor local neural survival. If, as we have argued, the polarity effect is more affected by local neural survival (specifically that of the peripheral processes) then this may have two implications for the potential improvement of patient outcomes by re-programming the clinical map. First, in devices such as the one studied here, where EMD variation across the array may be less than in the Cochlear CI24RE device, variations in spatial selectivity may be more strongly driven by neural survival, and it may be beneficial to have a measure that is specific to that parameter. Second, computational models (e.g. Goldwyn et al., 2010) indicate that, in devices that allow focused stimulation methods such as the tripolar mode, the appropriate intervention in cases of locally high thresholds depends on the cause. Specifically, Bierer and Litvak (2016) have argued that electrodes with a large EMD should be stimulated in tripolar mode, whereas those in neural “dead regions” should be de-activated. Again, a measure that differentiates between these two situations could provide significant clinical advantages.

## Summary and conclusions

i. Consistent with recent reports, polarity sensitivity at threshold varies across listeners and electrodes. Specifically, thresholds were lower for TP-C than for TP-A pulses for 27% of the electrode-subject combinations tested here.
ii. There was a significant trend for thresholds to be lower for more apical electrodes. However the polarity effect did not vary significantly as a function of longitudinal electrode position.
iii. There was a modest but significant across-electrode correlation between the polarity effect and the average threshold for the two polarities. The direction of the correlation was that lower thresholds for TP-C than for TP-A pulses corresponded to lower average thresholds. The correlation was in the same direction but larger than observed in two recent reports.
iv. Across subjects, the polarity effect correlated with the rate discrimination at high rates, as measured by Cosentino *et al* (Cosentino et al., 2016). The direction of the effect was that lower thresholds for TP-C than for TP-A pulses corresponded to better rate discrimination.
v. The results are consistent with, but do not prove, the idea that the polarity effect is an indicator of good neural survival, which in turn influences detection thresholds and performance on some supra-threshold tasks.

## Acknowledgments

We thank Lindsay DeVries for generously providing the EMD measurements from her study. This research was supported by awards RG91365 (Medical Research Council, Carlyon), MC-A060-5PQ75 (Action on Hearing Loss, Carlyon), and DC012142 (National Institute of Health, Arenberg).

